# Hierarchy and networks in the transcriptional response of *Mycobacterium abscessus* to antibiotics

**DOI:** 10.1101/2023.03.16.533064

**Authors:** Kelley Hurst-Hess, Charity McManaman, Yong Yang, Shamba Gupta, Pallavi Ghosh

## Abstract

*Mycobacterium abscessus* causes acute and chronic pulmonary infection in patients with chronic lung damage. It is intrinsically resistance to antibiotics effective against other pathogenic mycobacteria largely due to the drug-induced expression of genes that confer resistance. Induction of genes upon exposure to ribosome targeting antibiotics proceeds via WhiB7-dependent and -independent pathways. WhiB7 controls the expression of >100 genes, a few of which are known determinants of drug resistance. The function of the vast majority of genes within the regulon is unknown, but some conceivably encode additional mechanisms of resistance. Furthermore, the hierarchy of gene expression within the regulon, if any, is poorly understood. In the present work we have identified 56 WhiB7 binding sites using chromatin immunoprecipitation sequencing (CHIP-Seq) which accounts for the WhiB7-dependent upregulation of 70 genes, and find that *M. abscessus* WhiB7 functions exclusively as a transcriptional activator at promoters recognized by σ^A^/σ^B^ We have investigated the role of 18 WhiB7 regulated genes in drug resistance and demonstrated the role of MAB_1409c and MAB_4324c in aminoglycoside resistance. Further, we identify a σ^H^-dependent pathway in aminoglycoside and tigecycline resistance which is induced upon drug exposure and is further activated by WhiB7 demonstrating the existence of a crosstalk between components of the WhiB7-dependent and -independent circuits.

**Abstract Importance:** The induction of multiple genes that confer resistance to structurally diverse ribosome-targeting antibiotics is funneled through the induction of a single transcriptional activator, WhiB7, by antibiotic-stalled ribosomes. This poses a severe restriction in *M. abscessus* therapy as treatment with one ribosome-targeting antibiotic confers resistance to all other ribosome-targeting antibiotics. Here we uncover the intricacies of the WhiB7 regulatory circuit, identify three previously unknown determinants of aminoglycoside resistance and unveil a communication between WhiB7 dependent and independent components. This not only expands our understanding of the antibiotic resistance potential of *M. abscessus* but can also inform the development of much needed therapeutic options.

## Introduction

The genus *Mycobacterium* is comprised of obligate pathogens and environmental bacteria that are either saprophytes or opportunistic pathogens, and are all known to demonstrate varying levels of intrinsic antibiotic resistance that restricts therapeutic options against their infections (1–4). This innate resistance is attributed to a combination of a complex cell wall that poses a permeability barrier and the expression of chromosomally encoded effectors that confer resistance (1–4). Antibiotic resistance effectors include efflux pumps as well as enzymes that either modify/inactivate the target or the drug; these can be constitutively expressed, but are more frequently induced as a consequence of the transcriptional reprogramming that occurs upon drug exposure. Within the genus, *Mycobacterium abscessus* is an exceptionally drug-resistant, rapidly-growing, non-tuberculous mycobacterium (NTM) causing acute and chronic pulmonary disease in patients with underlying lung damage such as cystic fibrosis (CF) and COPD, as well as skin and soft tissue infections post-surgery/-trauma (5–8). The multi-drug resistance phenotype of *M. abscessus* presents a major challenge in treatment of its infections (7, 9). The current therapy involves a combination of a macrolide (clarithromycin or azithromycin) and intravenous amikacin and cefoxitin/imipenem for ~12-18 months with an average clearance rate of ~45% (10). These dismal eradication rates are attributed, at least in part, to antibiotic induced induction of resistance determinants that limit the efficacy of the regimen (11–13).

The best studied master-regulator of the antibiotic-induced reprogramming in mycobacteria is WhiB7 which belongs to the WhiB-like (Wbl) family of transcriptional regulators found exclusively in actinomycetes (14). It is characterized by the presence of four invariant cysteine residues that bind a [4Fe-4S] cluster, a conserved β-turn G[I/V/L]W[G/A]G) motif present in all Wbl proteins, a WhiB7 specific (EPW) motif adjacent to the β-turn and a C-terminal DNA binding AT-hook (RGRP) (14, 15). *whiB7* is one of the earliest genes induced to varying extents when mycobacteria are exposed to sub-inhibitory concentrations of structurally unrelated antibiotics that target both the 50S and 30S subunits of the ribosome; these include macrolides, lincosamides, ketolides, oxazolidinones, chloramphenicol, tetracyclines, aminoglycosides, phenicols and pleuromutilins (12, 15–17). A deletion of *whiB7* in *M. abscessus, M. smegmatis* and *M. tuberculosis* results in hypersensitivity to most, but not all, antibiotics that induce its expression (12, 17, 18). Most prominently, subinhibitory levels of tetracycline (TET) strongly induces *whiB7* in *M. abscessus*, but expression of the TET resistance determinant is WhiB7 independent (18). In addition to antibiotics, *whiB7* expression is induced by compounds that perturb respiration and redox balance, and, physiological stresses such as heat shock, iron starvation and entry into stationary phase (15–17, 19, 20). Finally, *whiB7* induction has also been observed upon infection of mycobacteria into macrophages and in lungs of infected mice suggesting its importance in virulence (21, 22).

The WhiB7 genes in actinobacteria are typically preceded by long leader sequences that include an upstream ORF (uORF) and a rho-independent terminator that function as an attenuator (23, 24). Ribosome stalling during starvation or in the presence of antibiotics leads to formation of an anti-terminator hairpin utilizing overlapping sequences in the uORF and the terminator leading to suppression of termination and induction of the downstream *whiB7* gene (23, 24). The WhiB7 protein then functions as a transcriptional activator and regulates the expression of >100 genes that comprise the WhiB7 regulon (12, 17). The function of a few genes within this regulon, including *erm41, eis2, hflX* and ABCFs, have been described in mycobacterial antibiotic resistance (12, 17, 25–28). However the role of the vast majority of genes within the regulon, as well as the hierarchy and the molecular networks, if any, are poorly understood in mycobacteria. In the present work we have determined the direct targets of *M. abscessus* WhiB7 using chromatin immunoprecipitation sequencing (CHIP-Seq) and demonstrated a role of MAB_1409c and MAB_4324c in aminoglycoside resistance. Additionally we also identify a σ^H^-dependent network involved in aminoglycoside and tigecycline resistance that communicates with the WhiB7 dependent pathway.

## RESULTS

### Identification of the WhiB7-dependent regulon of *M. abscessus*

In a previous study we compared the changes in gene expression of Δ*MabwhiB7* and Δ*MabwhiB7::phspMabwhiB7* strains in the absence of antibiotic using RNA sequencing ( RNAseq) and identified 229 genes comprising the *whiB7* responsive regulon in *M. abscessus* using the criteria of >2-fold induction in the *whiB7* overexpressing strain as compared to the Δ*MabwhiB7* mutant (*p_adj_* <0.01) (12). To identify genes that are dependent upon WhiB7 for expression, we followed the global changes in gene expression in wild-type *M. abscessus* and a Δ*MabwhiB7* strain upon antibiotic exposure. The elucidation of gene expression changes in Δ*MabwhiB7* was facilitated by the fact that TET strongly induces *whiB7* expression; however Δ*MabwhiB7* is not TET susceptible since the determinants of TET resistance lie outside the WhiB7 regulon. This enabled us to determine the changes in gene expression in wild-type and Δ*MabwhiB7* backgrounds under identical antibiotic exposure conditions. Upon treatment of the strains with 16μg/ml TET (1XMIC) for 45 mins, we identified 181 genes that were differentially regulated >2-fold *(p_adj_* <0.05). Of these, the expression of 170 genes was reduced >2-fold in Δ*MabwhiB7* compared to wild-type bacteria, while the expression of 11 genes increased >2-fold in the mutant. (Figure 1a-b, Supplementary data).

**Figure 1.**
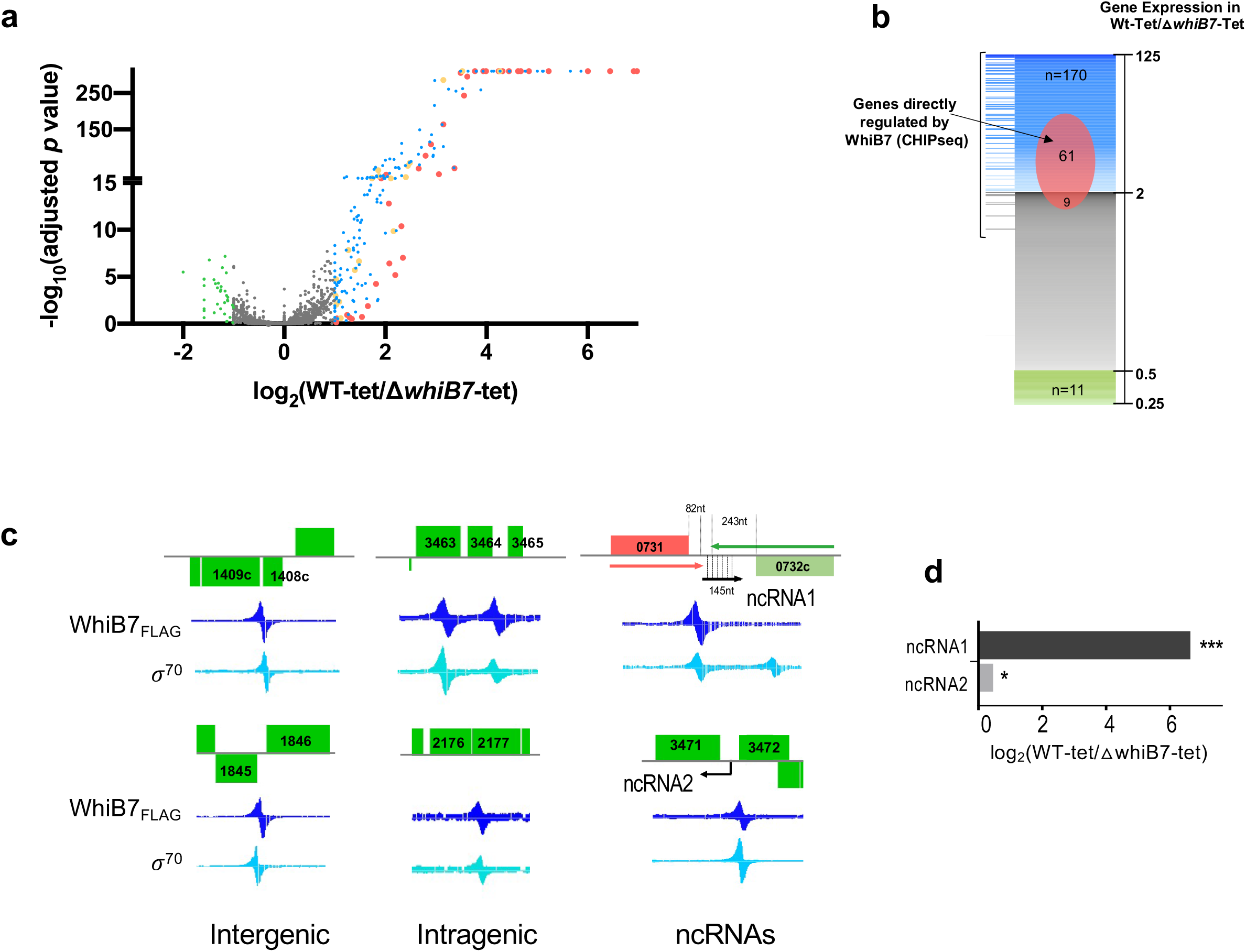
Determination of the *M. abscessus* WhiB7 dependent regulon and WhiB7 binding sites. **a)** Volcano plot of differentially expressed genes in *M. abscessus* ATCC19977 wild-type and Δ*MabwhiB7* strains upon exposure to TET (16μg/mL for 45 mins) determined by RNASeq. Four biological replicates were used of each sample. Genes differentially upregulated >2-fold in wild-type are indicated in blue, red and orange, and genes differentially downregulated >2-fold in wild-type are indicated in green. The red circles correspond to genes immediately adjacent to WhiB7 binding sites and orange circles correspond to operonic genes. **b)** Graphical representation of the WhiB7 regulon showing proportion of genes identified using CHIPSeq. Location of direct targets of WhiB7 shows the fold downregulation of each target in Δ*whiB7* **c)** Representative examples of WhiB7 binding sites at intergenic locations, within genes and transcribing ncRNAs visualized on Signal Map are shown. **d)** WhiB7 dependent induction of ncRNAs in wild-type and Δ*MabwhiB7* strains upon exposure to TET (16μg/mL for 45 mins). Region of complementarity between ncRNA1 and MAB_0732c transcript is shown using dashed vertical lines. *p_adj_=* 0 (***) and <0.01 (*).

### MabWhiB7 directly regulates the expression of a subset of 64 genes within its regulon

The 181 genes comprising the MabWhiB7 regulon presumably include both direct and indirect targets of WhiB7. To identify the genes that are directly regulated by WhiB7 we determined the genome-wide binding profile of MabWhiB7 using CHIP-Seq. For this purpose we constructed 3 strains: i) the *MabwhiB7* gene was 3X-FLAG tagged at the C-terminus, integrated at the L5 attachment site of a Δ*MabwhiB7* strain and expressed using its native promoter and upstream regulatory elements contained within a 650 nt leader sequence (Δ*whiB7::p_nat_whiB7_C-FLAG_*), ii) the Mab*whiB7* gene was 3X-FLAG tagged at the C-terminus, integrated at the L5 attachment site of a Δ*MabwhiB7* strain and constitutively expressed (Δ*whiB7::p_hsp_whiB7_C-FLAG_*) iii) Mab*whiB7* gene was 3X-FLAG tagged at the N-terminus and constitutively expressed from a chromosomal location of a Δ*MabwhiB7* strain (Δ*whiB7::p_hsp_whiB7_N-FLAG_*). Control experiments showed that addition of a C-terminal or N-terminal FLAG-tag did not interfere with WhiB7 function. Nonetheless maximum complementation was observed when Mab*whiB7* was expressed from its native promoter; we therefore used this strain for all CHIP-Seq experiments (Figure S1). The strain Δ*whiB7::p_nat_whiB7_C-FLAG_* was grown to mid-exponential phase (OD_600_= 0.6) followed by treatment with 16μg/ml TET for either 45 mins or 2hrs; DNA-nucleoprotein complexes were immunoprecipitated with anti-FLAG or anti-σ^70^ monoclonal antibodies followed by library preparation and Illumina sequencing. Using a previously published Python script, Peakcaller (29), we identified a total of 56 WhiB7 binding sites (Table 1) of which, 51 were located between genes that were transcribed either divergently or in the same direction; 5 peaks were found to be intragenic (Figure 1c,Table 1). Furthermore, all WhiB7 binding sites were also associated with a σ^70^ peak that recognizes both Group I and II sigma factors, implying that WhiB7 can affect transcription exclusively at promoters transcribed by σ^A^/σ^B^ (30) (Table 2).

**Table 1.** CHIPseq of MabWhiB7 binding sites. Genetic location and coordinates of peaks, gene function and fold downregulation in Δ*MabwhiB7* (RNAseq) are shown. Identity of gene regulated by WhiB7 binding from MEME analysis is also included. Genes in red correspond to peaks detected at 2hs of induction only.

| Peak # | WhiB7 CHIPseq peak | | | Gene Regulated | Gene description | Fold downregulation in ΔwhiB7-Tet (RNAseq) |
| --- | --- | --- | --- | --- | --- | --- |
|  | Coordinates | FAT score | Genetic location |  |  |  |
| 1 | 3842377 | 1 | MAB_3786c/ MAB_3787 | MAB_3786c | Hypothetical protein | 125.79 |
| 2 | 2411990 | 2 | MAB_2355c/MAB_2356 | MAB_2355c | Putative ABC transporter ATP-binding protein | 120.42 |
| 3 | 4615555 | 2 | MAB_4532c/MAB_4533 | MAB_4532c | Acetyltransferase | 86.89 |
| 4 | 3011060 | 1 | upstream of MAB_2956 | MAB_2956 | hypothetical protein - GATB Yqey superfamily | 64.05 |
| 5 | 1136821 | 1 | upstream of MAB_1125c | MAB_1125c | GNAT acetyltransferase | 38.44 |
| 6 | 4404776 | 1 | upstream of MAB_4324c | MAB_4324c | Putative GNAT acetyltransferase, RimI superfamily | 37.53 |
| 7 | 4370213 | 1 | upstream of MAB_4294 | MAB_4294 | Probable aspartate aminotransferase | 28.58 |
| 8 | 158505 | 1 | MAB_0163c/MAB_0164 | MAB_0163c | Aminoglycoside phosphotransferase | 25.62 |
| 9 | 3095974 | 1 | upstream of MAB_3042c | MAB_3042c | HflX | 24.65 |
| 10 | 2893879 | 1 | MAB_2844c/MAB_2845 | MAB_2845 | aconitate methyltransferase | 21.84 |
| 11 | 2954769 | 2 | upstream of MAB_2903 | MAB_2903 | Hypothetical protein | 21.77 |
| 12 | 2828542 | 1 | upstream of MAB_2780c | MAB_2780c | MFS transporter | 19.90 |
| 13 | 1346473 | 1 | MAB_1344c/MAB_1345 | MAB_1344c | Putative glucose-4,6-dehydratase/ oxidoreductase | 18.80 |
| 14 | 2345935 | 1 | upstream of MAB_2297 | MAB_2297 | erm41 | 15.90 |
| 15 | 1413707 | 11 | upstream of MAB_1409c | MAB_1409c | Putative drug antiporter protein precursor (MFS) | 15.30 |
| 16 | 4707211 | 1 | upstream of MAB_4621c | MAB_4621c | Putative GNAT acetyltransferase, Rim L superfamily | 13.64 |
| 17 | 3966090 | 1 | upstream of MAB_3913 | MAB_3913 | Putative translocator (RhtB family- efflux?) | 12.27 |
| 18 | 1299192 | 1 | MAB_1295c/MAB_1296 | MAB_1296 | Hypothetical protein | 11.75 |
| 19 | 1341921 | 1 | upstream of MAB_1340 | MAB_1340 | nucleotide binding protein (YgdF family) | 11.23 |
| 20 | 3507827 | 4 | upstream of MAB_3465 | MAB_3465 | Putative sulfate transporter/antisigma-factor antagonist | 8.33 |
| 21 | 874984 | 1 | upstream of MAB_0880 | MAB_0880 | MFS transporter | 6.94 |
| 22 | 405480 | 33 | upstream of MAB_0404c | MAB_0404c | Putative GNAT acetyltransferase | 6.31 |
| 23 | 2944991 | 1 | upstream of MAB_2892c | MAB_2892c | PdxS pyridoxal biosynthesis lyase | 5.08 |
| 24 | 1844715 | 11 | MAB_1845c/MAB_1846 | MAB_1846 | Putative ABC transporter ATP-binding protein | 4.57 |
| 25 | 5030041 | 1 | MAB_4921c/MAB_4922 | MAB_4921c | probable integral membrane protein | 4.04 |
| 26 | 2712704 | 1 | MAB_2670c/MAB_2671 | MAB_2670c | Histidinol dehydrogenase(AAB) | 4.99 |
| 27 | 4932689 | 1 | upstream of MAB_4820 | MAB_4820 | Hypothetical protein | 4.84 |
| 28 | 3737457 | 2 | upstream of MAB_3683c | MAB_3683c | Tryptophanyl-tRNA synthetase | 4.18 |
| 29 | 1984334 | 1 | upstream of MAB_1987 | MAB_1987 | phosphoheptulonate synthase | 4.22 |
| 30 | 3588702 | 4 | MAB_3543c/MAB_3544 | MAB_3544 | amino-acyl tRNA deacylase | 3.77 |
|  |  |  |  | MAB_3543c | RNA polymerase sigmaH | 2.45 |
| 31 | 1632207 | 1 | upstream of MAB_1603 | MAB_1603 | Valyl-tRNA synthetase | 3.52 |
| 32 | 340031 | 1 | MAB_0342c/MAB_0343 | MAB_0343 | Aspartate kinase | 3.15 |
| 33 | 2357379 | 1 | upstream of MAB_2305c | MAB_2305c | Hypothetical protein | 3.13 |
| 34 | 1129660 | 2 | MAB_1118c/MAB_1119 | MAB_1118c | Hypothetical protein | 2.90 |
| 35 | 3551647 | 59 | upstream of MAB_3509c | MAB_3509c | Hypothetical protein (uORF of whiB7) | 2.52 |
| 36 | 3130952 | 1 | upstream of MAB_3084c | MAB_3084c | tetrahydrodipicolinate synthase (AAB) | 2.37 |
| 37 | 3821725 | 2 | upstream of MAB_3763 | MAB_3763 | cutinase | 2.17 |
| 38 | 3012118 | 1 | MAB_2957/MAB_2958 | MAB_2958 | Putative transport protein | 2.06 |
| 39 | 3470985 | 3 | upstream of MAB_3424c | MAB_3424c | 2- isopropyl malate synthase | 2.04 |
| 40 | 3820662 | 13 | MAB_3762/MAB_3761c | MAB_3762 | permease: transporter | 1.92 |
| 41 | 3760436 | 1 | upstream of MAB_3705 | MAB_3705 | Tet family regulator | 1.85 |
| 42 | 2689063 | 1 | upstream of MAB_2647c | MAB_2647c | Anthranilate synthase | 1.77 |
| 43 | 1216748 | 1 | upstream of MAB_1201c | MAB_1201c | GreA | 1.53 |
| 44 | 3252647 | 2 | upstream of MAB_3211c | MAB_3211c | carbonic anhydrase | 1.43 |
| 45 | 4204628 | 1 | MAB_4139/ MAB_4138c | MAB_4139 | putative transcriptional regulator, ArsR family | 1.30 |
| 46 | 3172940 | 1 | upstream of MAB_3131c | MAB_3131c | Initiation factor IF-2 | 1.26 |
| 47 | 2382586 | 3 | MAB_2329c/MAB_2330 | MAB_2329c | hypothetical protein, MspA like porin | 1.12 |
| 48 | 2089239 | 1 | Lys tRNA/MAB_2089 | MAB_2089 | Hypothetical protein | 1.02 |
| Intragenic Peaks |  |  |  |  |  |  |
| 49 | 3507096 | 4 | inside MAB_3463 | MAB_3464 | Hypothetical protein | 8.86 |
| 50 | 3077676 | 1 | inside MAB_3023c | MAB_3022 ? | Hypothetical protein | 1.03 |
| 51 | 2191174 | 1 | N-terminal of MAB_2177 | MAB_2177 | probable ABC transporter, periplasmic | 7.47 |
| 52 | 3670771 | 1 | N-terminal of MAB_3621 | MAB_3621 | Probable taurine dioxygenase | 1.77 |
| 53 | 4941059 | 2 | inside MAB_4827 | unknown | n/a | n/a |
| Probable RNAs |  |  |  |  |  |  |
| 54 | 3513806 | 4 | upstream of MAB_3472 | MAB_3471 | Opposite orientation of motif; antisense RNA detected | ncRNA:1.4<br>MAB_3471: 1.4 |
| 55 | 736028 | 9 | MAB_0731/ MAB_0732c | MAB_0732 | Wrong direction for both ORFs; antisense RNA detected | ncRNA:99<br>MAB_0732c: 10.3 |
| 56 | 3252670 | 1 | upstream of MAB_3211c | opposite direction | Wrong orientation for MAB_3211c; antisense to MAB_3212c ? |  |

**Table 2.** Coordinates of peaks recognized by anti-σ^70^ antibodies (corresponding to σA and σ^B^ binding), location of TSS sites when available and spacing between −35 of WhiB7 MEME and TSS/-10.

| SigA CHIP-seq peak | WhiB7 CHIPseq peak | Gene Regulated | TSS | Spacing between motif and TSS | Spacing between motif and -10 |
| --- | --- | --- | --- | --- | --- |
| Coordinates | Coordinates |  |  |  |  |
| 3842419 | 3842377 | MAB_3786c | 3842397 | 33 | 24 |
| 2411983 | 2411990 | MAB_2355c | 2411993 | 34 | 24 |
| 4615553 | 4615555 | MAB_4532c | 4615566 | 34 | 24 |
| 3011065 | 3011060 | MAB_2956 | na | na | 23 |
| 4404754 | 4404776 | MAB_4324c | 4404783 | 34 | 24 |
| 3095971 | 3095974 | MAB_3042c | 3095977 | 35 | 24 |
| 2954778 | 2954769 | MAB_2903 | 2954778 | 33 | 23 |
| 2828533 | 2828542 | MAB_2780c | 2828544 | 33 | 24 |
| 2345951 | 2345935 | MAB_2297 | 2345927 | 22 | 13 |
| 1413694 | 1413707 | MAB_1409c | 1413705 | 33 | 24 |
| 4707208 | 4707211 | MAB_4621c | 4707223 | 32 | 23 |
| 3966094 | 3966090 | MAB_3913 | na | na | 24 |
| 1341898 | 1341921 | MAB_1340 | 1341922 | 24 | 14 |
| 3507836 | 3507827 | MAB_3465 | 3507835 | 32 | 23 |
| 874942 | 874984 | MAB_0880 | na | na | 22 |
| 405478 | 405480 | MAB_0404c | na | na | 23 |
| 2944981 | 2944991 | MAB_2892c | na | na | 14 |
| 1844720 | 1844715 | MAB_1846 | 1844710 | 33 | 23 |
| 5030047 | 5030041 | MAB_4921c | 5030054 | 34 | 24 |
| 3588682 | 3588702 | MAB_3544 | 3588768 | 25 | 14 |
|  |  | MAB_3543c | 3588704 | 21 | 13 |
| 1129635 | 1129660 | MAB_1118c | 1129677 | na | 16 |
| 3551617 | 3551647 | MAB_3509c | 3551635 | 33 | 23 |
| 3470974 | 3470985 | MAB_3424c | 3470992 | 34 | 23 |
| 3820679 | 3820662 | MAB_3762 | 3820266 | na | 24 |
| 1216740 | 1216748 | MAB_1201c | 1216746 | 31 | 22 |
| 4204594 | 4204628 | MAB_4139 | na | na | 18 |
| 2383570 | 2382586 | MAB_2329c | na | na | 23 |
| 2089231 | 2089239 | MAB_2089 | 2089297 | 33 | 23 |
| 3012165 | 3012118 | MAB_2958 | 3012152 | 32 | 23 |
| 340040 | 340031 | MAB_0343 | na | na | 13 |
| 3172950 | 3172940 | MAB_3131c | 3172964 | 23 | 13 |
| 4370260 | 4370213 | MAB_4294 | 4370226 | 23 | 13 |
| 158510 | 158505 | MAB_0163c | 158473 | 71 | 61 |
| 1299265 | 1299192 | MAB_1296 | 1299233 | 33 | 23 |
| 1346465 | 1346473 | MAB_1334c | 1346466 | 22 | 13 |
| 1632240 | 1632207 | MAB_1603 | na | na | 27 |
| 1984375 | 1984334 | MAB_1987 | 1984351 | 23 | 14 |
| 2689050 | 2689063 | MAB_2647c | 2689049 | 23 | 14 |
| 2712715 | 2712704 | MAB_2670c | na | na | 14 |
| 3130970 | 3130952 | MAB_3084c | 3130981 | 23 | 14 |
| 3737460 | 3737457 | MAB_3683 | 3737464 | 23 | 14 |
| 3670772 | 3670771 | MAB_3621c | 3670784 | 22 | 13 |
| 2191192 | 2191174 | MAB_2177 | 2191189 | 34 | 24 |
| 2357392 | 2357379 | MAB_2305c | 2357398 | 34 | 24 |
| 2893879 | 2893879 | MAB_2845 | 2893900 | 33 | 23 |
| 4932661 | 4932689 | MAB_4820 | 4932688 | 33 | 23 |
| 1136824 | 1136821 | MAB_1125c | 1136840 | 35 | 25 |
| 3821739 | 3821725 | MAB_3763 | 3821719 | 34 | 24 |
| 3252664 | 3252647 | MAB_3211c | na | na | 23 |
| 3760426 | 3760436 | MAB_3705 | 3760427 | 24 | 14 |
| **Probable RNAs** |  |  |  |  |  |
| 3513782 | 3513806 | upstream of MAB_3472 | 3513798 | 30 | 22 |
| 736042 | 736028 | MAB_0731/ MAB_0732c | 736030 | 30 | 22 |
| 3252670 | 3252670 | upstream of MAB_3211c | n/a |  |  |

Next, we analyzed the sequences corresponding to the MabWhiB7 peaks using MEME tools and identified a conserved motif comprised of an AT-rich sequence 3bp upstream of the −35 promoter element in all 56 CHIP-Seq regions (Figures 2a and b). The orientation of the conserved WhiB7 binding motif was then used to determine the identity of the genes regulated by WhiB7 binding and are shown in Tables 1 and 2. Using this method, 48 of 51 intergenic WhiB7 peaks were found to potentially regulate 49 *M. abscessus genes*. We used the RNAseq dataset to determine if the 49 genes identified above were differentially expressed in the Δ*MabwhiB7* strain when exposed to TET. Table 1 shows that the expression of 44 genes is significantly (*p_adj_* <0.01) downregulated >1.5-fold in Δ*MabwhiB7*, whereas the expression of 5 genes remains unchanged. However, none of the genes directly regulated by WhiB7 showed an increase in expression in Δ*MabwhiB7* implying that WhiB7 functions exclusively as a transcriptional activator. We also investigated the 5 intragenic WhiB7 binding peaks closely as they were all associated with a σ^70^ peak. Of these the WhiB7 binding sites within MAB_2177 and MAB_3621 were located at the extreme N-terminal of the ORFs, suggesting that these genes are likely misannotated. The significance of WhiB7 binding within MAB_3023c, MAB_3463 and MAB_4827 is unknown but could be driving the expression of downstream genes MAB_3022c and MAB_3464. Indeed, we find the expression of MAB_2177 and MAB_3464 to be strongly WhiB7 dependent (Table 1).

**Figure 2.**
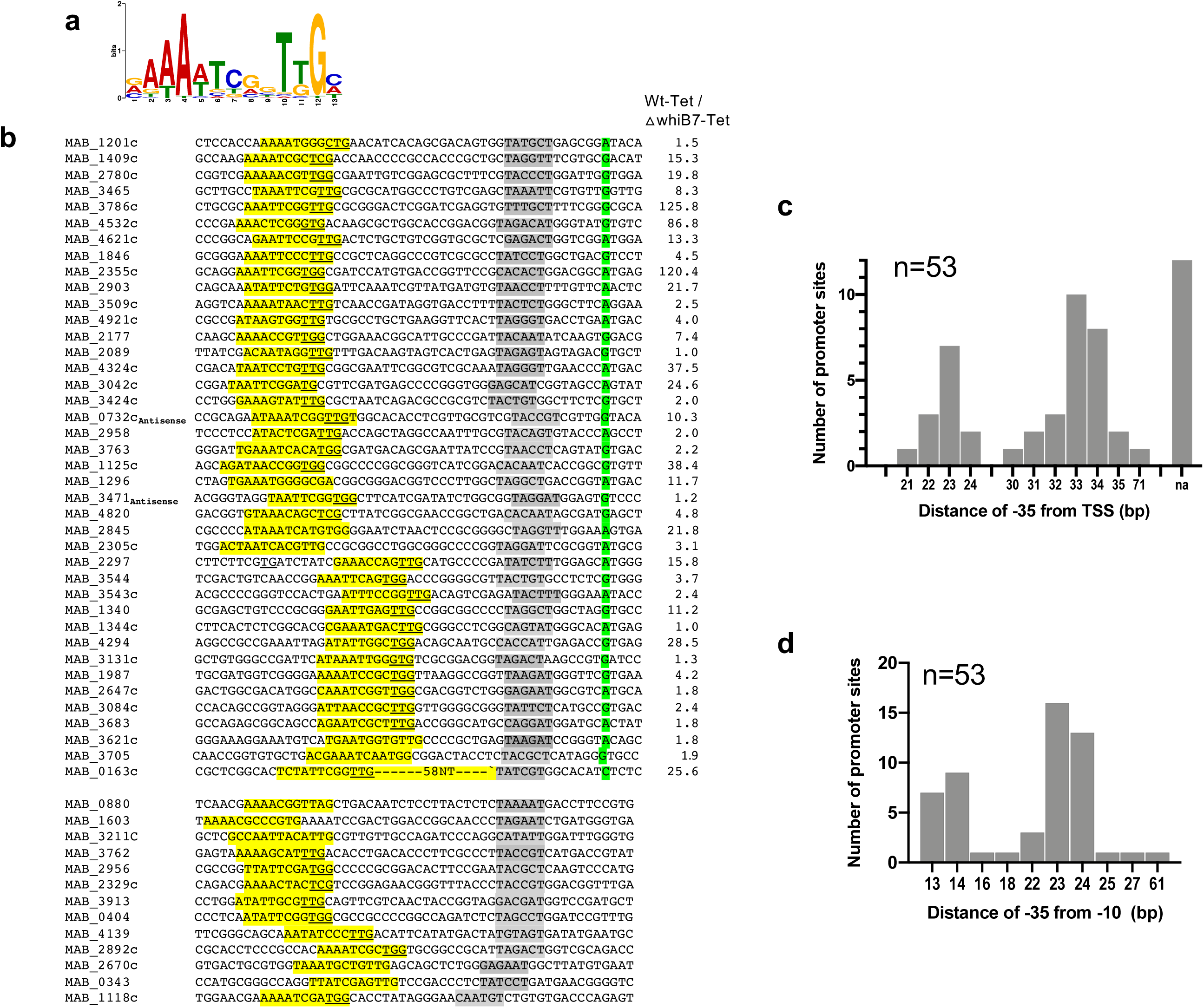
Conserved motif within WhiB7 binding sites. **a)** Sequence logo of enriched motif in WhiB7_FL_A_G_ bound sites identified using MEME Suite 5.5.1 (MEME E-value = 3.7.0e^-015^) showing the presence of an AT rich sequence separated by 3nts from a −35 site. **b)** Location of WhiB7 motif, known TSSs and potential −10 sequences within WhiB7_FL_A_G_ binding sites. Fold downregulation of each genes in Δ*whiB7* compared to wild-type strain is also noted. **c)** Histogram showing the frequency distribution of distances between the −35 sequence and known TSSs. **d)** Histogram showing the frequency distribution of distances between −35 sequence and potential −10 sequences corresponding to the σ^A^/σ^B^ consensus TANNNT.

Following identification of WhiB7 targets we mapped known transcription start sites (TSSs) with relation to the WhiB7 binding sites (31). Figure 2b-c and Table 2 show that most the −35 elements were located either 22-23nt or 33-34 nt from the TSS. A sequence corresponding to the −10 consensus sequence of σ^A^ /σ^B^ specific promoters (TANNNT) could also be identified either 13-14 nt or 23-24nt downstream of the −35 element and is consistent with previous studies in *M. tuberculosis* that demonstrate a cooperative binding between σA and WhiB7 (15, 30) (Figure 2b,d, Table 2). However we do not observe a correlation between the degree of dependence of WhiB7 in gene expression and the spacing between −35 and −10 elements.

Finally, we determined if the genes that are likely to be within the same transcription unit as the direct targets of WhiB7 were differentially expressed in wild-type *M. abscessus* when compared to Δ*MabwhiB7* (RNAseq). We identified 19 genes whose expression was WhiB7 dependent and were likely to be upregulated directly by WhiB7 binding upstream of the operon (Table 3). Therefore binding of WhiB7 to 56 chromosomal locations unequivocally accounted for the WhiB7 dependent upregulation of 70 genes (49 intergenic, 2 intragenic and 19 operonic genes).

**Table 3.** Expression of operonic genes regulated by WhiB7.

| Gene immediately downstream of WhiB7 binding site | Fold under-represented in $\Delta whiB7$ mutant | Operonic genes | Function | Fold under represented in $\Delta whiB7$ mutant |
| --- | --- | --- | --- | --- |
| MAB_3786c | 125.79 | MAB_3785c | Lipoprotein LppF | 19.21 |
| MAB_1846 | 4.57 | MAB_1847 | Metalloprotease | 5.44 |
|  |  | MAB_1848 | Hypothetical protein | 3.63 |
| MAB_2297 | 15.90 | MAB_2298 | Dephospho-CoA kinase | 4.47 |
| MAB_2892c | 5.08 | MAB_2890c | Glutamine amidotransferase PdxT | 5.59 |
|  |  | MAB_2891c | acyl-CoA thioesterase II | 5.29 |
| MAB_3763 | 2.17 | MAB_3764 | epoxide hydrolase EphA | 2.02 |
| MAB_2780c | 19.90 | MAB_2779c | Glyceraldehyde-3-phosphate dehydrogenase | 2.04 |
|  |  | MAB_2778c | phosphoglycerate kinase | 3.33 |
| MAB_1340 | 11.23 | MAB_1341 | hypothetical protein | 8.86 |
| MAB_0880 | 6.94 | MAB_0881 | tRNA/rRNA methyltransferase | 3.67 |
| MAB_2958 | 2.05 | MAB_2959 | Putative N-acetyltransferase, Rim I related | 2.03 |
| MAB_2647c | 1.76 | MAB_2646c | Membrane protein in trp biosynthesis | 2.41 |
| MAB_3084 | 2.37 | MAB_3083 | ribonuclease J | 3.33 |
| MAB_3683c | 4.19 | MAB_3682c | membrane protein | 2.13 |
| MAB_1603 | 3.52 | MAB_1604 | folylpolyglutamate synthase FolC | 11.50 |
| MAB_0343 | 3.15 | MAB_0344 | 50S ribosomal protein L28 | 4.31 |
|  |  | MAB_0345 | Hypothetical protein | 2.79 |
| MAB_3131c | 1.26 | MAB_3130 | Ribosome-binding factor A (RbfA) | 2.63 |

### Regulation of non-coding RNAs by MabWhiB7

The WhiB7 motif at 3 locations were oriented in a direction opposite to that of the ORFs within the region and are suggestive of driving the expression of ncRNAs (Figure 1c, Table 1). From the RNAseq dataset we detected two ncRNAs in the region between MAB_0731 and MAB_0732c (ncRNA1) and between MAB_3471 and MAB_3472 (ncRNA2) consistent with a previous report (31). The expression of ncRNA1 increases upon treatment with TET and is downregulated ~100-fold (*p_adj_* <0.001) in the Δ*MabwhiB7* mutant (Figure 1d, Supplementary data). Interestingly, ncRNA1 is complementary along most of its length to the 3’-UTR of MAB_0732c (Figure 1c). Since the expression of MAB_0732c is downregulated ~10-fold in Δ*MabwhiB7* it is likely that ncRNA1 functions by stabilizing MAB_0732c mRNA and merits further investigation. A differential change in expression of ncRNA2 was not observed in the Δ*MabwhiB7* mutant and its function is not immediately obvious.

### Characterization of direct targets of WhiB7 identifies novel effectors of drug resistance

The direct targets of WhiB7 belong to a variety of functional categories including genes encoding ribosome associated proteins, transcription regulators, acetyltransferases, transporters, biosynthetic and metabolic enzymes and hypothetical proteins. Of these, only 7 genes have been characterized previously and have known functions in *M. abscessus* drug resistance (Figure 3a, Table 4) ; the role of the majority of genes regulated by WhiB7 (either directly or indirectly) is unknown. We therefore used a complementation assay in which a gene under investigation was constitutively expressed from a chromosomal location of a Δ*MabwhiB7* strain, followed by analysis of its ability to restore the susceptibility of Δ*MabwhiB7* to that of wild-type bacteria. Recently we used this assay to identify the role of MAB_2780c in spectinomycin resistance (32). A spectrum of 8 drugs that target both the 50S and 30S ribosomal subunits were evaluated-amikacin (AMK), tigecycline (TIG), streptomycin (STR), spectinomycin (SPC), apramycin (APR), erythromycin (ERT), clarithromycin (CLA) and clindamycin (CLIN). We tested 16 genes belonging to different functional categories-acetyltransferases, putative transporters and hypothetical proteins. As seen in Figure 3b, MAB-1409c, a putative MFS efflux pump, restores APR sensitivity of Δ*MabwhiB7* to a level exceeding that of wild-type *M. abscessus*. Further, MAB_1409c is also moderately active in complementing the AMK and STR sensitivity of Δ*MabwhiB7* indicating that it could function in the efflux of a variety of aminoglycoside antibiotics. Additionally, MAB_4324c, a GNAT N-acetyltransferase, also confers moderate levels of resistance to AMK alone (Figure 3b, S2a-d). No apparent role for the remaining 14 genes could be detected in resistance to the tested antibiotics (Figure S2a-d).

**Figure 3.**
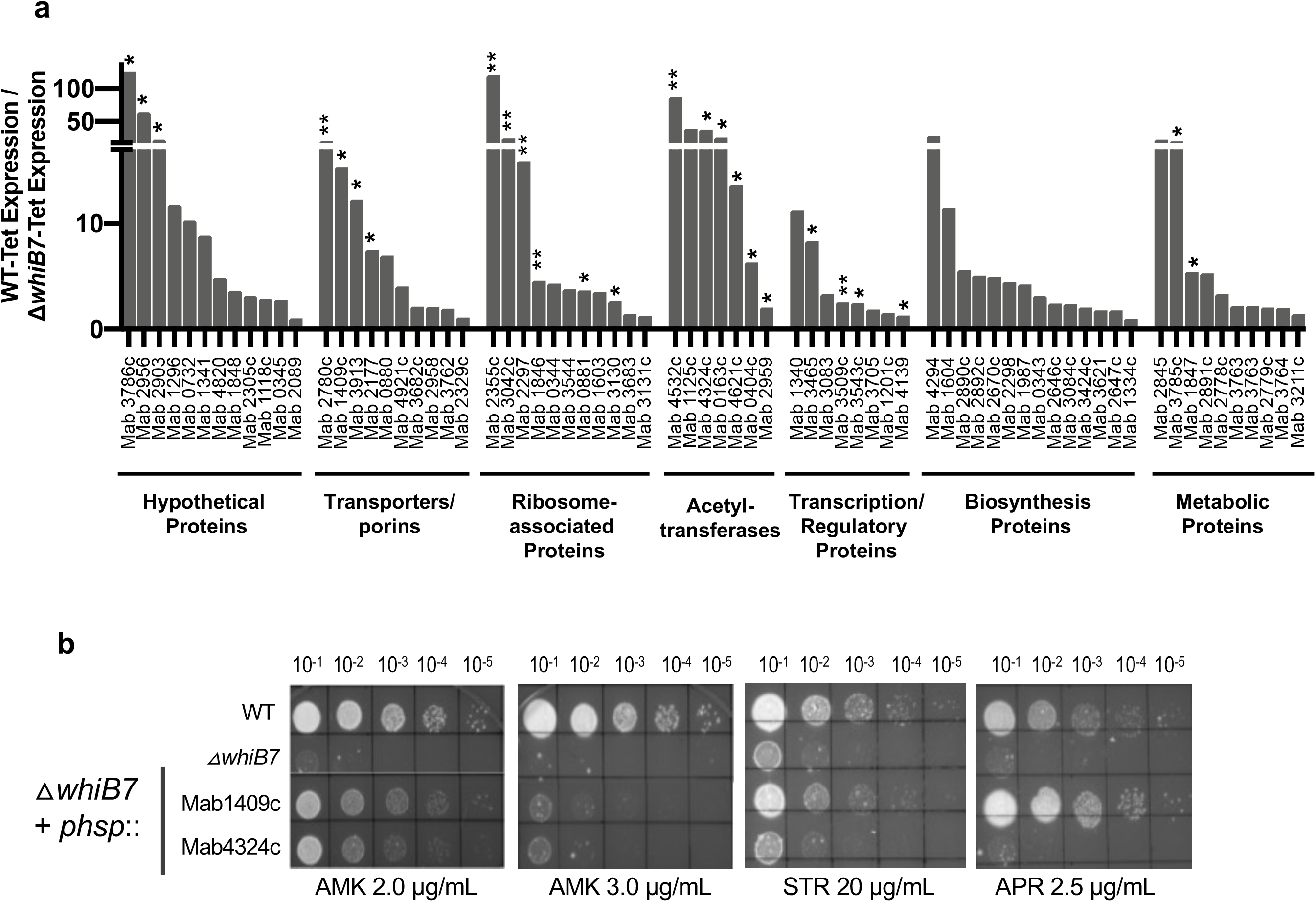
Functional analysis of WhiB7 regulated genes. **a)** Functional classification of genes directly regulated by WhiB7 and the fold down-regulation in a Δ*MabwhiB7* is shown. Genes with known roles in drug resistance is indicated with **. Genes that have been evaluated in this study are indicated with *. **b)** Growth of ten-fold serial dilutions of *M. abscessus* ATCC 19977, Δ*MabwhiB7*, and Δ*MabwhiB7* complemented with either MAB_1409c or MAB_4532c on Middlebrook 7H10 OADC containing indicated concentrations of AMK, STR and APR. Data is representative of >3 independent experiments.

**Table 4.** Genes within the direct regulon of WhiB7 with known functions and tested in this study.

| Gene Regulated | Gene description | Role in Drug resistance |
| --- | --- | --- |
| MAB_3509c | Hypothetical protein (uORF of whiB7) | Expression of whiB7 (22) |
| MAB_1846 | Putative ABC transporter ATP-binding protein | Lincosamide (L) (24) |
| MAB_2297 | erm41 | MLSKO (34) |
| MAB_2355c | Putative ABC transporter ATP-binding protein | Macrolide (M) (27) |
| MAB_2780c | MFS transporter | SPC (31) |
| MAB_3042c | HflX | Macrolide/lincosamide (25) |
| MAB_3543c | RNA polymerase sigmaH factor | TIG (32) |
| MAB_4532c | Acetyltransferase | AMK (26) |
| MAB_1409c | Putative drug antiporter protein precursor (MFS) | **APR, STR, AMK** |
| MAB_3543c | RNA polymerase sigma-E factor | **AMK, STR, SPC, APR** |
| MAB_4324c | Putative acetyltransferase, GNAT | **AMK** |
| MAB_0163c | Aminoglycoside phosphotransferase | Not detected |
| MAB_0404c | Putative GNAT acetyltransferase | Not detected |
| MAB_0881 | tRNA/rRNA methyltransferase | Not detected |
| MAB_1847 | Metalloprotease | Not detected |
| MAB_2177 | probable ABC transporter, periplasmic | Not detected |
| MAB_3785c | Lipoprotein LppF | Not detected |
| MAB_2903 | Hypothetical protein | Not detected |
| MAB_2956 | hypothetical protein - GATB Yqey superfamily | Not detected |
| MAB_2959 | Putative N-acetyltransferase, Rim I related | Not detected |
| MAB_3130 | Ribosome-binding factor A (RbfA) | Not detected |
| MAB_3465 | Putative sulfate transporter/antisigma-factor antagonist | Not detected |
| MAB_3786c | Hypothetical protein | Not detected |
| MAB_3913 | Putative translocator (RhtB family- efflux?) | Not detected |
| MAB_4139 | putative transcriptional regulator, ArsR family | Not detected |
| MAB_4621c | Putative acetyltransferase | Not detected |

### SigH requlates pathways required for resistance to TIG, AMK, APR and STRP

The CHIP-Seq data set shows that WhiB7 directly affects expression of 70 genes within the WhiB7 regulon of 181 genes. This suggests a hierarchical control of gene expression in which WhiB7 directly affects transcription of 70 genes while the remaining genes are indirectly WhiB7 dependent and are expressed by one or more transcriptional regulators contained within the direct targets of WhiB7. We therefore tested three transcriptional regulators that are direct targets of WhiB7-MAB_3543c (*sigH*), MAB_3465 (a putative anti-sigma factor antagonist) and MAB_4139 (ArsR) for their ability to complement the drug sensitivity of Δ*MabwhiB7*. While MAB_4139 and MAB_3465 did not display a role in antibiotic resistance, expression of *sigH* effectively restored the STR and APR sensitivity of Δ*MabwhiB7* wo wild-type levels, and moderately complemented the AMK and TIG sensitivity of Δ*MabwhiB7* (Figure 4a, Figure S2b,d); its effect on 50S targeting antibiotics was negligible (Figure S3). To study the role of *sigH* in intrinsic antibiotic resistance of *M. abscessus*, we constructed an isogenic deletion of *sigH* (MAB_3543c) in *M. abscessus* ATCC 19977. The Δ*MabsigH* strain was mildly sensitive to AMK and TIG in comparison to Δ*MabwhiB7*, suggesting a modest contribution of *sigH* to AMK and TIG resistance (Figure 4c-d). In contrast, Δ*MabsigH* and Δ*MabwhiB7* strains displayed comparable sensitivity to STR and APR suggesting a significant involvement of *sigH* in STR and APR resistance (Figure 4c). Furthermore, overexpression of the regulator of σ^H^, *rshA*, in wild-type bacteria recapitulated the drug sensitivity of Δ*MabsigH*, a phenotype compatible with increased levels of RshA sequestering σ^H^ leading to antibiotic sensitivity (Figure 4b-c). Our results are consistent with previous reports which demonstrate that point mutations in *sigH* and its anti-sigma factor, *rshA*, are associated with increased TIG resistance (33, 34) and additionally uncover an involvement of *σ*^H^ in aminoglycoside resistance.

**Figure 4.**
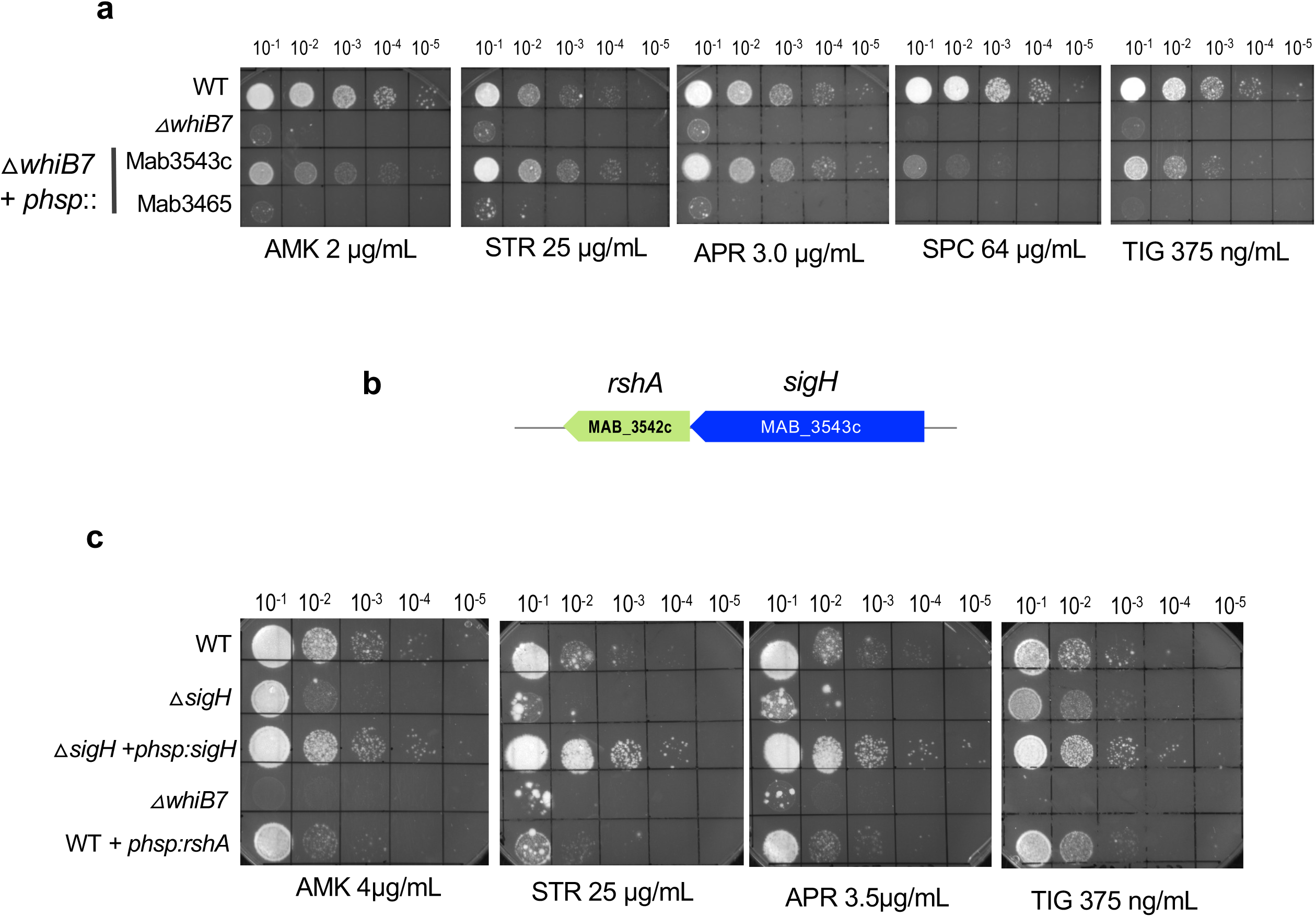
Functional analysis of transcriptional regulators within the direct regulon of WhiB7. **a)** Growth of ten-fold serial dilutions of *M. abscessus* ATCC 19977, Δ*MabwhiB7*, and Δ*MabwhiB7* complemented with either MAB_3453c or MAB_3465 on Middlebrook 7H10 OADC containing indicated concentrations of AMK, STR, TIG, SPC and APR. Data is representative of >3 independent experiments. **b)** Genomic organization of *sigH* (MAB_3543c) and its anti-sigma factor *rshA* (MAB_3542c). **c)** Growth of ten-fold serial dilutions of *M. abscessus* ATCC 19977, Δ*MabwhiB7*, Δ*MabsigH*, Δ*MabsigH* complemented with *sigH* and wild-type *M. abscessus* overexpressing *rshA* on Middlebrook 7H10 OADC containing indicated concentrations of AMK, STR, TIG, and APR. Data is representative of >3 independent experiments.

These results are suggestive of a scenario in which expression of *sigH* is activated by WhiB7, which is then required for expression of a subset of genes within the WhiB7 regulon. We therefore determined the global changes in gene expression in a Δ*MabsigH* strain treated with sublethal doses of TET and compared it to a wild-type strain. We identified 33 genes (not including *sigH*) that were differentially regulated > 2-fold (*p_adj_* <0.01) (Figure 5a-b, Figure S5, Supplementary data) and expected that these would include genes within the WhiB7 regulon. Surprisingly, deletion of *sigH* did not significantly impact (>2-fold; *p_adj_* <0.01) the expression of any of the genes within the WhiB7 regulon implying that the indirect targets of WhiB7 are not dependent on *σ*^H^ (Figure 5b-d, Supplementary data). Furthermore, the expression of the 33 *σ*^H^ dependent genes were largely unchanged in a Δ*MabwhiB7* strain (Figure 5e). These results were confounding since overexpression of *sigH* in Δ*MabwhiB7* either partially, or significantly, complemented the drug sensitivity of Δ*MabwhiB7* (Figure 4a). A closer inspection revealed that expression of *sigH* is ~12-fold induced by TET in wild-type bacteria but is also induced ~ 5-fold in Δ*MabwhiB7;* the expression of *sigH* is therefore activated only 2-fold in the presence of WhiB7(Figure 5f). Since the σ^H^ regulon is unchanged in the Δ*MabwhiB7* strain, it suggests that the levels of functional σ^H^, which is additionally controlled at the post-translational level by anti-σ^H^, does not change significantly in the Δ*MabwhiB7* strain (Figure 7). Nonetheless, overexpression of *sigH* in the drug hypersusceptible Δ*MabwhiB7* background can modestly enhance the drug tolerance of Δ*MabwhiB7* and implies that a subset of genes within the σ^H^ regulon can play a role in resistance to TIG and aminoglycoside antibiotics.

**Figure 5.**
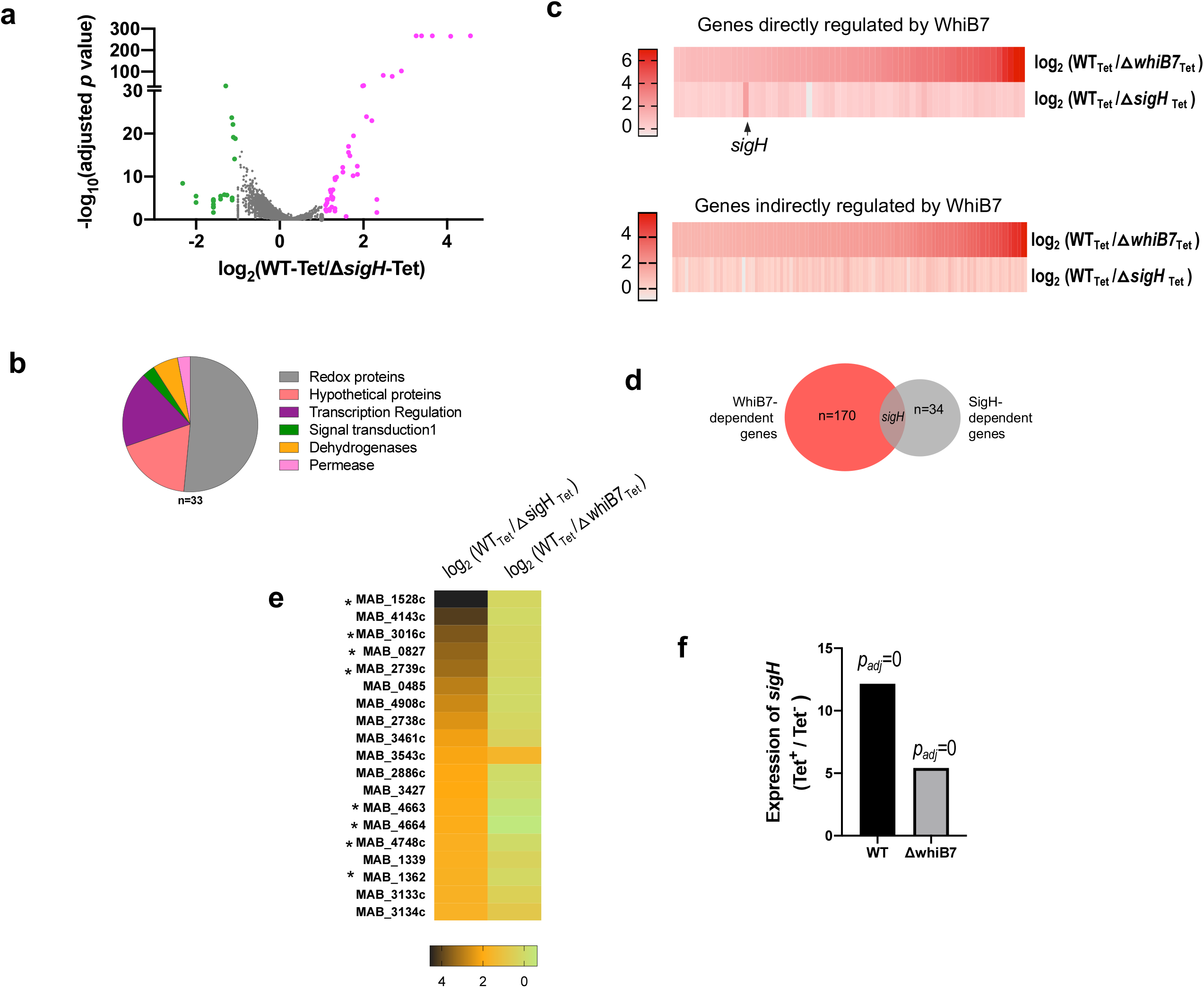
Evaluation of the role of *sigH* in *M. abscessus* drug resistance. **a)** Volcano plot of differentially expressed genes in *M. abscessus* ATCC19977 wild-type and Δ*MabsigH* strains upon exposure to TET (16μg/mL for 45 mins) determined by RNASeq. Four biological replicates were used of each sample. Genes differentially upregulated >2-fold in wild-type are indicated in purple, and genes downregulated >2-fold in wild-type are indicated in green. **b)** Pie chart of functional categories of σ^H^ regulated genes. **c)** Heat map showing differential expression of WhiB7 regulon genes in Δ*MabsigH* and Δ*MabwhiB7* strains exposed to TET (16μg/mL for 45 mins). **d)** Venn diagram showing overlap of the WhiB7 and σ^H^ regulons. **e)** Heat map showing differential expression of the σ^H^ regulon genes (downregulated >2-fold in Δ*MabsigH)* in Δ*MabsigH and ΔMabwhiB7* backgrounds upon exposure to TET (16μg/mL for 45 mins). **f)** Expression of *sigH* in wild-type and Δ*MabwhiB7* strains upon exposure to TET (16μg/mL for 45 mins) using RNAseq.

### MAB_4664 and MAB_1362 within the σ^H^ regulon mediate drug resistance

About half of the σ^H^ regulon is represented by proteins involved in redox pathways; the remaining belong to diverse functional categories ( Figure 5b). To determine which of the 33 σ^H^ dependent genes mediate drug tolerance, we evaluated the function of 8 genes belonging to different functional categories – oxidoreductases (MAB_1528c, MAB_ 3016c, MAB_0827, MAB_2739c, MAB_4748c), hypothetical proteins (MAB_4663, MAB_4664) and a regulatory protein (MAB_1362/*sigE*). Figure S4 shows that none of the oxidoreductases tested could complement the drug-sensitive phenotype of Δ*MabsigH*. Overexpression of MAB_4664, a gene encoding a hypothetical protein with no apparent sequence homologue, was found to consistently restore the TIG sensitivity of both Δ*MabsigH* and Δ*MabwhiB7* to wild-type levels (Figure 6a and b). In addition, overexpression of MAB_*s/gE* in Δ*MabsigH* and Δ*MabwhiB7* backgrounds greatly increased the tolerance of the mutant strains to STR, AMK and APR but not to TIG. This suggests a hierarchical control of gene expression within the σ^H^ regulon, in which MAB_4664 expression is independent of σ^E^ and is controlled by σ^H^ alone, whereas σ^E^ controls expression of genes within the σ^H^ regulon that confer AMK, STR and APR tolerance (Figure S5c). An analysis of the upstream regions of σ^H^ dependent genes using MEME tools displayed the presence of a canonical σ^H^/σ^E^ motif previously described in *M. tuberculosis* (Figure S5a and b) (35, 36). Curiously we note a variability in the nucleotide immediately downstream of the conserved −35 GGAA motif within the σ^H^ regulated genes; MAB_4664 and MAB_1362 which appear to be direct targets of σ^H^, contain a G residue whereas the remaining genes with an T/A/C at that location are likely to be regulated by σ^E^ and/or σ^H^ (Figure S5b).

**Figure 6.**
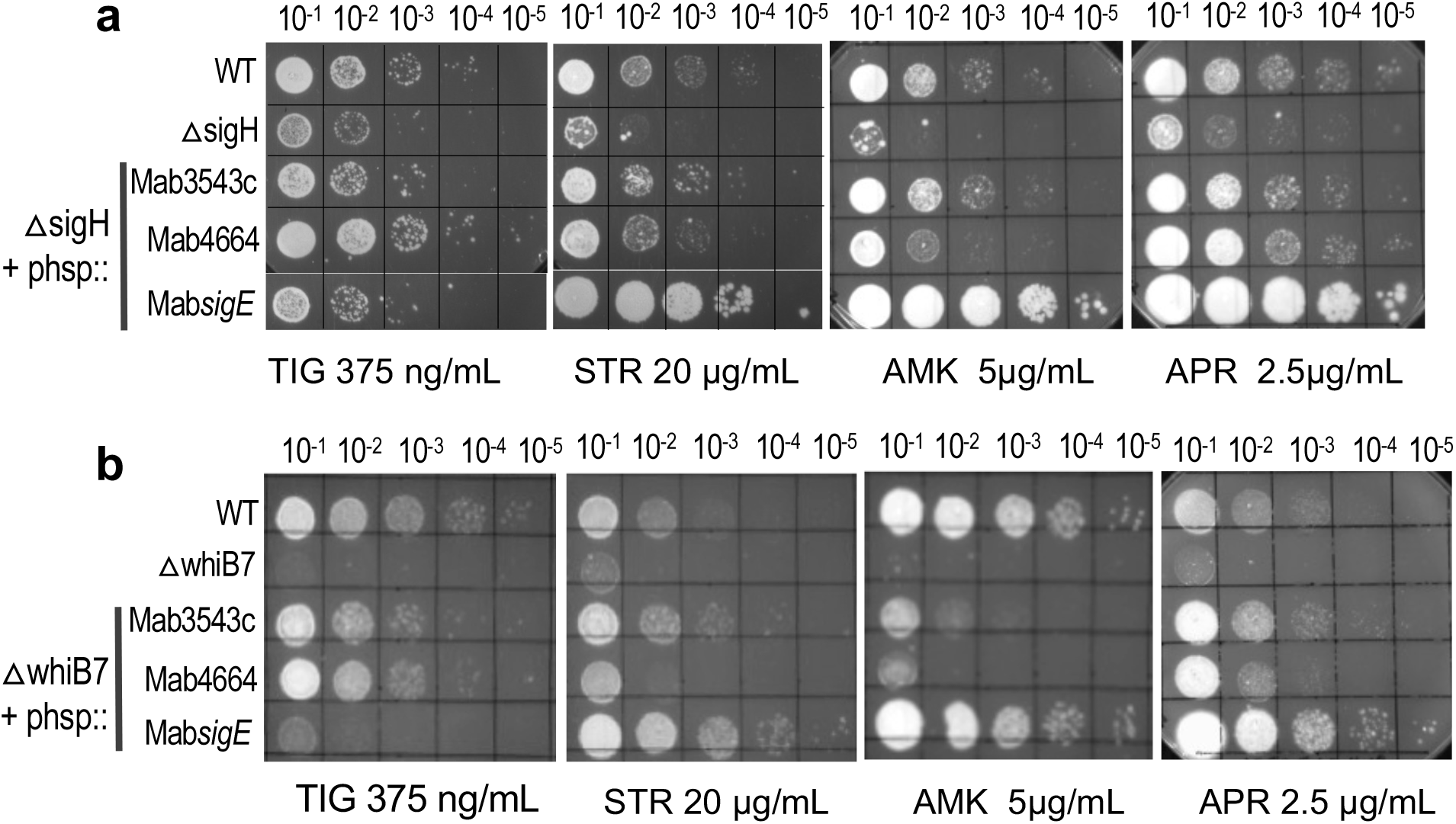
The *sigH* dependent genes, MAB_4664 and MAB_1362, confer drug resistance. **a)** Growth of 10-fold serial dilutions of *M. abscessus* ATCC 19977, Δ*MabsigH* and Δ*MabsigH* complemented with *sigH, MAB_4664 and MAB_1362 (sigE)* on Middlebrook 7H10 OADC containing indicated concentrations of AMK, STR, TIG, and APR. Data is representative of >3 independent experiments. **b)** Growth of 10-fold serial dilutions of *M. abscessus* ATCC 19977, Δ*MabwhiB7* and Δ*MabwhiB7* complemented with *sigH, MAB_4664 and MAB_1362 (sigE)* on Middlebrook 7H10 OADC containing indicated concentrations of AMK, STR, TIG, and APR. Data is representative of >3 independent experiments.

**Figure 7.**
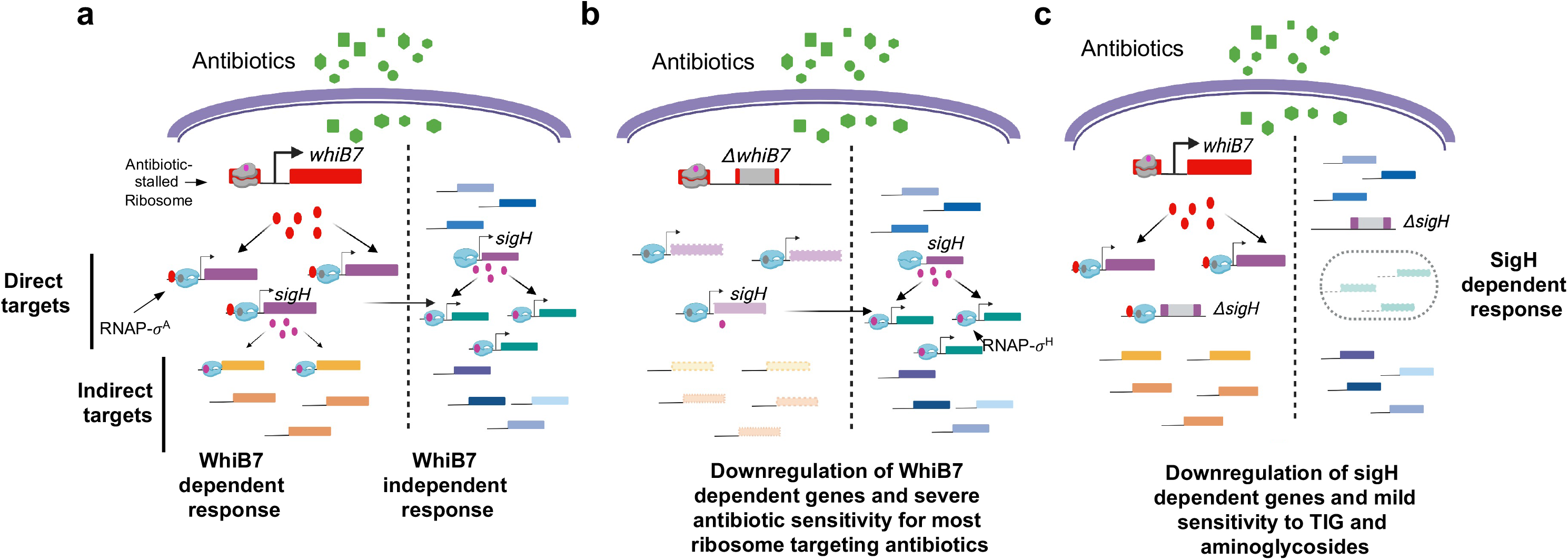
Model showing a cross-talk between WhiB7 and σ^H^ regulons. **(a) Global reprogramming of gene expression in wild type bacteria** by ribosome targeting antibiotics (green) occurs via WhiB7 dependent and independent pathways. WhiB7 regulated genes are shown in purple (direct targets) and orange (indirect targets). Genes induced independent of WhiB7 are shown in green and blue. Induction of *sigH* is antibiotic inducible and WhiB7 independent but is augmented 2-fold in the presence of WhiB7. (**b) Pattern of gene induction in a Δ*MabwhiB7* background.** Downregulated genes are shown in faded purple and orange shades. Induction of *sigH* and its regulon proceeds in Δ*MabwhiB7* despite the absence of WhiB7 activation. **(c) Pattern of gene induction in a Δ*MabsigH* background**. Deletion of *sigH* results in downregulation of the *sigH* regulon. The WhiB7 regulon remains unchanged.

## Discussion

Herein we demonstrate using CHIP-Seq that *M. abscessus* WhiB7 binds to 56 chromosomal locations. Using a combination of RNAseq and MEME analysis tools we then determine the identity of genes regulated by WhiB7 binding. Whilst the WhiB7 regulon of 181 genes comprises of 11 genes that are downregulated, the 70 genes that are direct targets of WhiB7 are all upregulated, confirming that WhiB7 functions exclusively as a transcriptional activator. The direct targets of WhiB7 comprise genes with diverse functions but are overrepresented by genes encoding transporters, acetyltransferases and ribosome associated proteins/proteins involved in amino-acid biosynthesis. Of the 70 direct targets of WhiB7, the function of only 7 have been clearly elucidated and the role of the remaining genes remain largely unknown (25–27, 32, 37, 38). We made use of a complementation assay to elucidate the role of 18 genes that included 5 putative acetyltransferases, 3 putative transporters, 3 transcription regulators, 3 hypothetical genes, 2 ribosome associated proteins and 2 metabolic enzymes, all of which were significantly drug inducible.

Acetyltransferases are known to transfer an acetyl group to diverse substrates from small molecules such as aminoglycosides and mycothiol to macromolecules (39). Aminoglycosides, classified as 2-DOS (APR, AMK, KAN) and non-DOS antibiotics (STR) based on their chemical structure, target slightly different portions of the 16S rRNA within the 30S ribosomal subunit (40). Resistance to aminoglycosides is commonly mediated by drug modification by acetyltransferases, phosphotransferases and nucleotidyltransferases (40). In *M. abscessus*, MAB_4395, an N-acetyltransferase (AAC2) is known to confer resistance to kanamycin B, tobramycin and dibekacin whereas MAB_4532c, an *eis* acetyltransferase confers AMK, but not APR or STR resistance (27). MAB_2385, a phosphotransferase outside the WhiB7 regulon, is the only known determinant of STR resistance (37). The *M. abscessus* genome encodes a large number of additional acetyltransferases and phosphotransferases several of which are included within the WhiB7 regulon and 6 are direct targets. We evaluated 3 out of 5 acetyltransferases with unknown functions. MAB_4324c, a putative GNAT acetyltransferase, showed a modest ability to confer AMK resistance but was not effective against STR, APR or SPC. Interestingly, MAB_4324c shows homology to the RimI superfamily of proteins that acetylate the S18 ribosomal proteins; the role of ribosome acetylation is hitherto unknown. Further work will be necessary to determine if AMK resistance by MAB_4324c is a consequence of acetylation of the antibiotic or the S18 ribosomal protein. MAB_0404, a GNAT acetyltransferase and MAB_4621c, a Rim L superfamily GNAT acetyltransferase however did not display an involvement in resistance to any of the antibiotics tested. This was surprising since MAB_0404c is strongly induced by ribosome targeting antibiotics. It is possible that MAB_0404c and MAB_4621c target antibiotics other than those tested in our study or catalyze acetylation of targets involved in pathways other than drug resistance.

The *M. abscessus* genome also encodes a large number of putative transporters and efflux pumps, several of which are within the WhiB7 regulon and are inducible by ribosome targeting antibiotics; 10 were found to be direct targets of WhiB7. However, the mere induction of a transporter does not confirm its role in drug resistance and the functions of the vast majority of efflux pumps are unknown. Recently we showed that MAB_2780c, a direct target of WhiB7, encodes a MFS efflux pump and is a major determinant of high-level SPC resistance in *M. abscessus* (32). An evaluation of 3 additional putative transporters – MAB_1409c, MAB_2177 and MAB_3913 was performed in this study of which only MAB_1409c demonstrated a significant ability to complement the APR sensitivity of Δ*MabwhiB7*. A mild complementation of AMK and STR sensitivity of Δ*MabwhiB7* by MAB_1409c was also observed and is consistent with a previous study (11); the role of MAB_1409c in APR resistance was however unexplored. Future characterization of MAB_1409c will enable determination of its substrate specificity and mechanisms of efflux.

WhiB7 directly regulates the expression of about half of the genes within its regulon which implies that a transcriptional regulator directly induced by WhiB7 can subsequently activate the next tier of genes within the regulon. Surprisingly, there were few transcriptional regulators that were directly regulated by WhiB7. MAB_3465, an anti-anti sigma factor, although a potentially interesting candidate, neither restored the drug sensitivity of Δ*MabwhiB7*, nor resulted in a global change in gene expression when expressed in wild-type bacteria (not shown). The ability of σ^H^ to complement TIG and aminoglycoside sensitivity of Δ*MabwhiB7* suggested that a subset of genes within the WhiB7 regulon were under the transcriptional control of σ^H^ Contrary to this expectation, RNAseq data demonstrated that the expression of the WhiB7 regulon was σ^H^ independent. σ^H^ was instead required for transcription of 33 genes outside the WhiB7 regulon. Why then did overexpression of σ^H^ complement the drug sensitivity of Δ*MabwhiB7?* We note that complementation by σ^H^ was only observed at lower concentrations of AMK, STR and TIG and that a Δ*MabsigH* strain was only mildly sensitive to the above antibiotics suggesting that σ^H^ plays a minor role in *M. abscessus* drug tolerance. Our results imply that some genes within the σ^H^ regulon confer low levels of tolerance to AMK, STR and TIG and that the effect of these genes are revealed when overexpressed in a hypersusceptible Δ*MabwhiB7* background. One such gene is MAB_4664 whose mechanism of action is unknown; the presence of a predicted transmembrane domain and involvement in tolerance to multiple antibiotics is suggestive of a role in efflux. In addition, a large majority of the σ^H^ regulated genes are involved in redox-related pathways consistent with a previous definition of the regulon using a *M. abscessus* strain containing a mutant *rshA*, as well as with the σ^H^ regulon of *M. tuberculosis* (35, 41). Coincidentally, several antibiotics, including aminoglycosides, have been shown to induce redox-related changes as a secondary aspect of their lethality (42). It is conceivable that redox proteins encoded within the *σ*^H^ regulon can mitigate this ancillary effect of antibiotics; consequently *σ*^H^ exerts a modest influence on antibiotic tolerance.

The activation of *sigH* expression by WhiB7 appears to be conserved in actinomycetes as the *Streptomyces coelicolor sigR* (the *sigH* homologue) is similarly under WhiB7_Sc_ control (43). However curiously the spectrum of resistance conferred by σ^H^ and σ^R^ in the two genera is different and implies the induction of distinct genes. Furthermore, CHIP-Seq analysis of WhiB7 in Streptomyces revealed the presence of 830 binding sites regulating at least 312 genes, which is vastly different from that observed in *M. abscessus* despite the presence of significant overlaps (44). While these could be attributed to differences in methods used to immunoprecipitate WhiB7, it is possible that the WhiB7 regulon is indeed different in these bacteria. Finally, it remains to be determined how the expression of over half of the WhiB7 regulon is induced in *M. abscessus*. While it is possible that some WhiB7 binding peaks have escaped detection in our experiments, it is an unlikely explanation since we have been unable to detect additional peaks in CHIP-Seq experiments utilizing two different locations of the FLAG tag and also expression of WhiB7 from constitutive promoters (not shown). Curiously we observe a striking absence of σ^A^ binding sites upstream of most of these indirect targets of WhiB7 which may suggest the involvement of an alternate sigma-factor that is post-transcriptionally activated in a WhiB7 dependent manner. Additionally, several WhiB7 target genes tested here do not associate with antibiotic resistance phenotypes. This may reflect a limitation in the assay where i) an insufficient number of antibiotics have been tested and ii) multigene interactions were not evaluated. Alternatively, the genes within the WhiB7 regulon may be involved in pathways other than drug resistance which is supported by previous observations of *whiB7* induction in *M. tuberculosis* under heat shock and low iron conditions and are presently under investigation (20).

## Materials and Methods

### Media and Bacterial Strains

*M. abscessus* ATCC19977 strains were grown at 37°C with shaking at 220 rpm in Middlebrook 7H9 (DIFCO) supplemented with 0.05% Tween 80 and 10% OADC. Antibiotics were added to indicated concentrations where required. An isogenic deletion in *MAB_3543c (sigH)* was constructed using recombineering, followed by removal of the zeocin cassette by Cre-mediated recombination at *loxP* sites as described previously(12). Unmarked deletion mutants were reconfirmed using PCR followed by sequencing of the PCR product. Complementing strains were created by cloning a gene under investigation in pMH94 under the control of the hsp60 promoter followed by integration at the phage L5 *attB* site of an appropriate deletion strain. Mab_whiB7 with a FLAG tag at either the N- or C-terminal was cloned in pMH94 under the control of the hsp60 promoter. Mab_whiB7 with a FLAG tag at the C-terminal was also cloned in pMH94 with its native promoter and upstream regulatory elements contained within a 650 nt leader sequence. All bacterial strains are listed in Table S1.

### Antibiotic Sensitivity Assays

Wild type and mutant strains of *M. abscessus* were grown to an A_600_ of 0.7-0.8. Cells were tested for drug susceptibility by spotting a 10-fold serial dilution on Middlebrook 7H10 (DIFCO) plates containing the indicated concentration of antibiotics.

### RNA preparation and RNA-Seq analysis

Wild type *M. abscessus ATCC 19977*, Δ*MabwhiB7*, Δ*MabsigH* were grown to exponential phase (OD=0.6-0.7) in Middlebrook 7H9-OADC and exposed to 16μg/mL of tetracycline (TET) for 45 minutes. Total RNA was prepared using the Qiagen RNA preparation kit followed by DNAse I treatment. Unexposed samples were used as controls. Approximately 5 *μ*g total RNA samples were treated with the Ribo-Zero™ rRNA removal procedure (Illumina) to enrich for mRNA. Approximately 500 ng of RNA was used for library preparation using the NEBNext® Ultra II RNA kit and high throughput sequencing on the Illumina NextSeq platform. The sequence data was analyzed using the reference based analysis and default parameters on Rockhopper v2.03 in which the data is normalized by upper quartile normalization and transcript abundance is reported as RPKM. Differential gene expression is tested for each transcript and q-values are then reported that control the false discovery rate (45, 46). RNAseq experiments were performed >3 independent times.

### Chromatin immunoprecipitation sequencing (CHIP-Seq) and data analysis

ChIP-Seq was performed as previously described with minor modifications (47). The Δ*whiB7::p_nat_whiB7_C-FLAG_, ΔwhiB7::p_hsp_whiB7_C-FLAG_* and *ΔwhiB7::p_hsp_whiB7_N-FLAG_* strains were grown at 37°C in Middlebrook 7H9 broth with 0.05% Tween 80 and 10% OADC to an OD_600_ = 0.6. The Δ*whiB7::p_nat_whiB7_c-FLAG_* strain was additionally induced with 16μg/mL of tetracycline (TET) for 45 minutes. This was followed by cross-linking using 1% formaldehyde for 30 min with constant agitation and quenching with 250mM glycine. The cells were pelleted, washed with TBS buffer and resuspended in buffer 1 (20mM HEPES (pH 7.5), 50mM KCl, 0.5mM DTT, 10% glycerol) containing a protease inhibitor cocktail (Sigma). Cells were harvested and lysed using the CryoMill (Retsch) in buffer 1 followed by sonication for 30 mins at the high setting of the Bioruptor™ sonicator (Diagenoge). The DNA protein complex was immunoprecipitated with either anti-FLAG monoclonal antibody M2 (Sigma) or anti-σ^70^ antibody (BioLegend) for 18 hours at 4°C and processed as previously described (47). Each CHIP-Seq experiment was performed using two biological replicates. Genomic DNA libraries enriched for WhiB7 binding were prepared using the NEB Next® Ultra II Library Prep kit for Illumina® followed by sequencing on the Illumina platform (Wadsworth Center, sequencing core facility). Reads were aligned to the reference genome using the Bowtie2 and SamTools algorithms (48). Regions of enrichment were identified using a custom Python script as described previously (29, 30). Relative enrichment are reported as fold over threshold (FAT) score. The enriched regions were analyzed using MEME Suite 5.5.1 using the default parameters (49).

## Acknowledgements

We thank The Wadsworth Center’s Applied Genomics Technology Core for sequencing of RNA-Seq and CHIP-Seq libraries and the Media Core for preparation of media and buffers. PG is supported by NIH awards AI155473 and AI146774 and the Wadsworth Center.

## Author Contributions Section

**Charity McManaman: :** performed experiments and analyzed data. **Kelley Hurst-Hess:** performed experiments. **Shamba Gupta** : performed experiments. Yong Yang: performed experiments **Pallavi Ghosh**: Conceptualizing, Data analysis, Methodology, Writing, Reviewing.

**Supplementary Data : Differential gene expression in Wt, *AMabwhiB7 and AMabsigH* in *Mab* ATCC 19977 upon exposure to a sublethal dose of TET (16μg/ml for 45 mins).**

**Table S1 List of strains used in the study**

## Supplementary Figures

**Figure S1: Ability of WhiB7 tagged strains to complement antibiotic sensitivity of *AMabwhiB7*.** Growth of ten-fold serial dilutions of *M. abscessus* ATCC 19977, Δ*MabwhiB7*, and Δ*MabwhiB7* complemented with either untagged *whiB7, whiB7FLAG_C-term_*, or *whiB7FLAG_N-term_* expressed from a constitutive promoter or *whiB7FLAG_C-term_* expressed from a native promoter on Middlebrook 7H10 plates containing ERT (7*μ*g/mL). Data is representative of >3 independent experiments.

**Figure S2: (a-d)** Growth of ten-fold serial dilutions of *M. abscessus* ATCC 19977, Δ*MabwhiB7*, and Δ*MabwhiB7* complemented with indicated genes on Middlebrook 7H10 plates containing indicated concentrations of antibiotics. Data is representative of >3 independent experiments.

**Figure S3:** Growth of ten-fold serial dilutions of *M. abscessus* ATCC 19977, Δ*MabwhiB7*, and Δ*MabwhiB7* complemented with either MAB_3543c, MAB_3465 and empty vector control on Middlebrook 7H10 plates containing indicated concentrations of 50S targeting antibiotics. Data is representative of >3 independent experiments.

**Figure S4:** Growth of ten-fold serial dilutions of *M. abscessus* ATCC 19977, Δ*MabsigH*, and Δ*MabsigH* complemented with either MAB_3543c, MAB_0827, MAB_1528c, MAB_3016c, MAB_4663, MAB_4748c or MAB_2739c on Middlebrook 7H10 plates containing indicated concentrations of AMK, TIG and STR. Data is representative of >3 independent experiments.

**Figure S5: a)** Sequence logo of enriched motif in SigH-dependent genes identified using RNAseq (downregulated >3-fold; *p_adj_* < 0.01) using MEME Suite 5.5.1. **b)** Location of conserved motif in upstream regions of SigH-dependent genes. Fold downregulation of each genes in Δ*sigH* strain is also noted. Difference in base composition of nucleotide immediately downstream to conserved GGAA motif is indicated in green (G/C) and yellow ( A/T). **c)** Possible hierarchy within the σ^H^ regulon.

## Supplementary Datasets

1. Transcriptome of *M. abscessus* wild type, Δ*whiB7* and Δ*sigH* strains exposed to 16 μg/mL tetracycline for 45 minutes:

